# DNA2 and MSH2 activity collectively mediate chemically stabilized G4 for efficient telomere replication

**DOI:** 10.1101/2025.04.04.647332

**Authors:** Anthony Fernandez, Tingting Zhou, Steven Esworthy, Changxian Shen, Helen Liu, Jessica D. Hess, Hang Yuan, Nian Liu, Guojun Shi, Mian Zhou, Settapong Kosiyatrakul, Vikas Gaur, Joshua Sommers, Winfried Edelman, Guo-Min Li, Robert Brosh, Weihang Chai, Marietta Y W T Lee, Dong Zhang, Carl Schildkraut, Li Zheng, Binghui Shen

**Author notes:** To whom correspondence should be addressed: Li Zheng. These authors contributed equally to this work (Anthony Fernandez, Tingting Zhou).

## Abstract

G-quadruplexes (G4s) are widely existing stable DNA secondary structures in mammalian cells. A long-standing hypothesis is that timely resolution of G4s is needed for efficient and faithful DNA replication. *In vitro*, G4s may be unwound by helicases or alternatively resolved via DNA2 nuclease mediated G4 cleavage. However, little is known about the biological significance and regulatory mechanism of the DNA2-mediated G4 removal pathway. Here, we report that DNA2 deficiency or its chemical inhibition leads to a significant accumulation of G4s and stalled replication forks at telomeres, which is demonstrated by a high-resolution technology: Single molecular analysis of replicating DNA (SMARD). We further identify that the DNA repair complex MutSα (MSH2-MSH6) binds G4s and stimulates G4 resolution via DNA2-mediated G4 excision. MSH2 deficiency, like DNA2 deficiency or inhibition, causes G4 accumulation and defective telomere replication. Meanwhile, G4-stabilizing environmental compounds block G4 unwinding by helicases but not G4 cleavage by DNA2. Consequently, G4 stabilizers impair telomere replication and cause telomere instabilities, especially in cells deficient in DNA2 or MSH2.

---

Efficient and faithful DNA replication is of paramount importance to cell survival and proper proliferation ^1–6^. However, DNA replisomes frequently face challenges including endogenous factors and/or environmental DNA-damaging agents ^1, 7–10^. These stress conditions cause DNA replication forks to stall, and if not stabilized or repaired, they collapse, resulting in mutations and chromosomal rearrangements, cell cycle arrest, or cell death. Repetitive DNA sequences such as micro-satellites or mini-satellites in centromeres, telomeres, and other loci may potentially form non-β-form DNA structures, which are “difficult-to-replicate” (DTR) ^11–15^. They are major endogenous sources of replication stress. Of the non-B-form DNA structures, G-quadruplexes (G4s) can spontaneously form and are thermodynamically stable. The G-rich single-strand DNA (ssDNA) sequences adopt a conformation which consists of stacks of two or more G-quartets formed by four guanines via Hoogsteen base-pairing stabilized by a monovalent cation ^16–19^.

In mammalian cells, G4s are prevalent as an epigenetic structural and functional motif across the genome ^20, 21^. G4s, if not resolved prior to DNA replication, may frequently cause problems for DNA polymerases ^22, 23^. Different mechanisms exist in mammalian cells to resolve and/or clean up the G4s barrier for efficient DNA replication. DNA helicases such as FANCJ and BLM can unwind G4s ^24, 25^. In addition, the ssDNA binding protein RPA (RPA1, RPA2, RPA3) or CST (CTC1, STN1, TEN1) may possibly unfold G4 structures ^26–28^. It is known that unfolding or unwinding G4s typically requires a ssDNA tail on which to load the RPA, CST, or G4 helicases ^24, 25, 27^. These two mechanisms are crucial for resolving G4s under normal physiology. However, if cells are exposed to environmental chemical compounds (ECCs) that induce or stabilize G4s, helicase-driven G4s unwinding or RPA-mediated G4 melting are inhibited ^24, 25^. Several industrial ECCs, endogenous metabolites, or drugs, including TMPyP4, can bind to, stabilize, and inhibit the unfolding or unwinding of G4s ^24, 25^, and newer technologies have permitted discovery of a greater variety of G4 stabilizing compounds ^29^.

Previously, we showed that the bifunctional mammalian helicase/nuclease DNA2 cleaves G4s or other secondary structures and that such nucleolytic activity is important in promoting centromeric and telomeric DNA replication ^30, 31^. However, the biological significance of this DNA2-mediated G4 resolution pathway requires more investigation. The impact of G4-inducing/stabilizing ECCs on this pathway is unclear. Furthermore, because G4 structures are implicated as chromosomal structural elements and epigenetic motifs that regulate gene expression ^20, 21, 32, 33^. DNA2-mediated G4s excision must be tightly controlled. Previous studies have reported many G4-binding proteins ^34, 35^. The function of these G4 binding proteins in the regulation of G4 resolution for DNA replication remains unknown.

In this study, we detect G4s in mammalian cells using a G4-specific antibody to determine that DNA2 deficiency or chemical inhibition leads to a significant accumulation of G4s. Furthermore, DNA2 deficiency potentiates G4 stabilizing compounds such as PIPER and TMPyP4 to increase G4 levels in cells. Using the single molecular analysis of replicating DNA (SMARD) technique ^36^, we show that DNA2 deficiency and/or G4 stabilizing compounds result in replication defects in telomeres, which contain the most abundant G4-forming sequences in the genome. We further identify that the mismatch repair protein MSH2 is a G4 binding protein *in vitro* and in cells. MSH2 is a key activator of DNA2 nuclease activity to cleave G4s *in vitro* and clean up G4s for DNA replication in mammalian cells. MSH2 is a member of the mismatch repair pathway for genome duplication fidelity. It forms a complex with MSH3 or MSH6 to recognize DNA mismatches or abnormal DNA structures ^37^. We discover that MutSα (MSH2-MSH6) interacts with DNA2 and stimulates DNA2 to cleave G4s and facilitate Polδ-mediated DNA synthesis. Our result is consistent with the previous report that MutSα binds to G4*invitro*^38^. MSH2 deficiency, like DNA2 deficiency or inhibition, causes G4s accumulation and defective telomere replication. In addition, G4-inducing and -stabilizing compounds such as PIPER block G4 resolution by helicases and cause significant G4 accumulation, particularly in DNA2 and MSH2 mutant cells. Our studies suggest an important role of MSH2 in facilitating DNA2-mediated G4 removal, especially in the presence of G4 stabilizing compounds.

## Results

### DNA2 is important in G4 resolution via G4 excision repair for efficient telomere DNA replication

We previously showed that purified recombinant DNA2 cleaved G4-bearing DNA substrates, mimicking the intramolecular G4 structures ahead of or within the DNA replication fork ^31^. To determine if DNA2 is important for removing G4s in cells, we performed immunofluorescence (IF) co-staining of DNA2 and G4, using anti-DNA2 and an anti-G4 structure-specific antibody ^22^, and used an Airyscan Joint Deconvolution (jDCV) confocal microscope with the joint deconvolution algorithm to visualize DNA2 and G4 DNA foci at sub-diffraction-limit resolution. We observed that a few DNA2 foci co-localized with G4 foci (Fig. 1a). To further demonstrate if DNA2 catalyzes G4 excision repair, we reconstituted G4 excision repair in an assay using purified recombinant DNA2 and Polδ and synthetic oligo based G4 DNA substrates (Fig. 1b, Top). In the G4 excision repair assay, DNA2 cleavage of G4s results in a DNA gap. A DNA polymerase (e.g., Polδ or Polβ) then fills in the DNA gap with ^32^P-labeled dTTP and the other three deoxyribonucleotides, producing ^32^P-labeled gap-filled products and fully extended products. We found that DNA2 and DNA polymerases effectively repaired G4s, producing duplex DNA (extended products), but in the absence of DNA2, DNA polymerase did not produce any repair products (Fig. 1b, bottom). To test if DNA2 is important for G4 resolution in cells, we carried out G4 IF staining in wild-type (WT) mouse endothelial fibroblasts (MEFs) and DNA2^+/-^ MEFs, which we previously generated and used to study the function of DNA2 in G4 resolution and telomere replication. We also analyzed G4 levels in WT MEFs treated with the DNA2 inhibitor C5. We found that DNA2 haploinsufficiency or DNA2 inhibition by C5 greatly increased G4 levels (Fig. 1c, d).

**Fig. 1.**
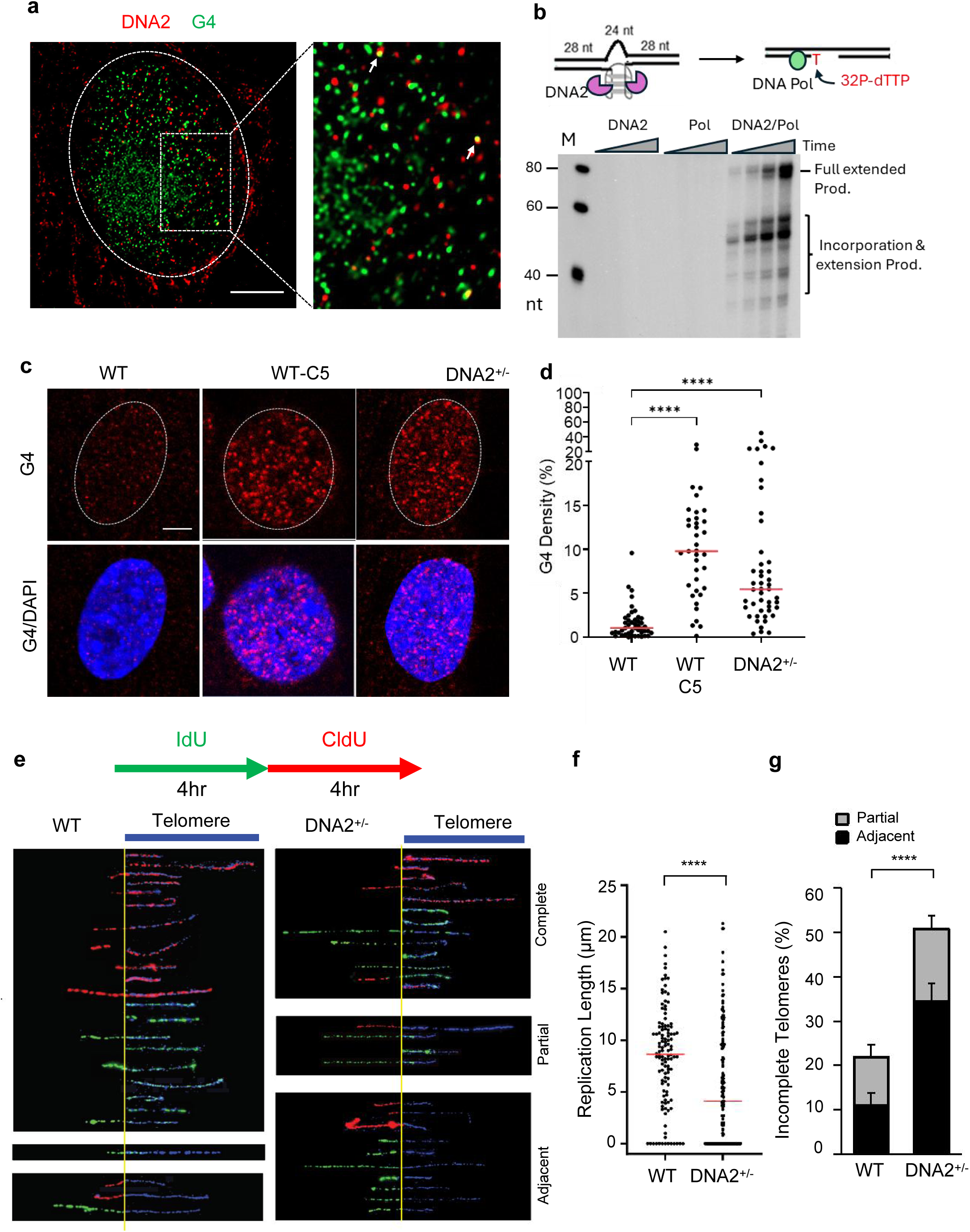
DNA2 deficiency leads to G4 accumulation in mammalian cells. **a** AIRYSCAN confocal microscopy of G4s; DNA2-G4 co-immunofluorescence staining in MEFs. The right panel shows the enlarged view of the boxed region. The spots with co-localized DNA2 and G4 foci were indicated with white arrows. Scale bar = 5 μm; **b** Reconstituted G4 excision repair assay. Top panel: schematic diagram elucidating biochemical assays for G4 excision, in which nucleases DNA2 cleave G4 to convert it into a gap. DNA polymerase such as Polβ, Polδ, or Polε then fills the gap using ^32^P-dTTP and the other three dNTP as substrates, producing ^32^P-labeled products. Bottom panel: Representative denaturing PAGE image showing G4 excision by DNA2 and the gap filling by Polβ; **c, d** G4 immunofluorescence staining in WT and DNA2^+/-^ MEFs. Panel **c** shows the representative AIRYSCAN confocal microscopy images of G4s, and Panel (**d**) shows the quantification of G4 in different cells. All p values were calculated using the Student’s *t*-test. * p < 0.05, ** p < 0.01, *** p < 0.001, and **** p < 0.0001. Scale bar = 5 μm; **e-g** DNA replication at telomeres in WT and DNA2^+/-^ MEFs; (**e**) Top: Scheme of the IdU (green) and CldU (red) pulse labeling. Bottom: SMARD microscopy images of replicated telomeres in WT and DNA2^+/-^ MEF cells. Imaged fibers contain both non-telomeric and telomeric DNA. SMARD fibers are arranged showing non-telomeric DNA at the left and telomeres on the right (indicated by the blue telomere drawing) aligned by start of the telomere at the yellow line. The top panels of SMARD fibers represent fully replicated telomeres with either IdU and/or CldU incorporation along the length of the telomere. Middle panels depict fibers with partial telomere replication, where IdU or CldU incorporated into some of the telomere but not for the entire length. The bottom panels depict completely un-replicated telomeres, where IdU and CldU incorporated immediately adjacent to, but not into telomeres. (**f**) Relative length of replicated telomeres in different cells. (**g**) Percentage of incompletely replicated telomeres in different cells. The frequency of partial telomere replication and replication fork stalling adjacent to telomeres were calculated.

To further determine the impact of G4 accumulation on DNA replication in cells, we analyzed replication specifically at the telomere region, which consists of the repetitive DNA sequence TTAGGG and has a high propensity for forming G4 structures ^39^. Using the SMARD approach to analyze replication at the individual DNA fiber levels, we analyzed replication dynamics exclusively in telomere sequences, using thymidine analogs CldU and IdU to label nascent DNA and a telomere-specific probe to define the telomere regions. We found in the WT MEFs within 8 hours that the mean length of replicated telomeres was 8.7 μm, while in DNA2^+/-^ MEFs it was 4.6 μm (Fig. 1e, f), exhibiting slower average replication fork speeds in DNA2^+/-^ MEFs. In addition, the frequency of incomplete telomere (partial and adjacent) increases from 22% (12% partial and 10% adjacent) in WT MEFs to 50% (35% partial and 15% adjacent) in DNA2^+/-^ MEFs (Fig. 1e, g). Consistently, inhibition of DNA2 by the DNA2 inhibitor C5 ^40^ in the WT cells caused similar replication stalling at telomeres (Supp. Fig.1). DNA2^-/-^ embryonic stem (ES) cells had significantly more telomere loss and sister telomere fusion than the WT ES cells (Supp. Fig. 2a-c).

### MutSα interacts with DNA2, stimulates its G4 cleavage activity, and facilitates telomere replication

We next explored the factors that contribute to DNA2-mediated repair of G4s. To this end, we expressed 3x-Flag-tagged DNA2 in 293T cells and carried out co-IP to pull down the 3x-Flag-tagged DNA2 (Fig. 2a) and its binding proteins. We used mass spectrometry to identify the DNA2-binding proteins. Our analysis revealed DNA replication and repair platform proteins such as RPA, RFC, and PCNA, which have previously been found in complex with DNA2 ^41–43^ (Fig. 2b), as well as several new DNA2 binding partners, including MMR proteins MSH2 and MSH6; nucleotide excision repair (NER) proteins RAD23A and RAD23B; NHEJ proteins Ku70/Ku80 and DNA-PK; and homology-directed repair (HDR) proteins RPA1, RPA2, and PARP1 (Fig. 2b).

**Fig. 2.**
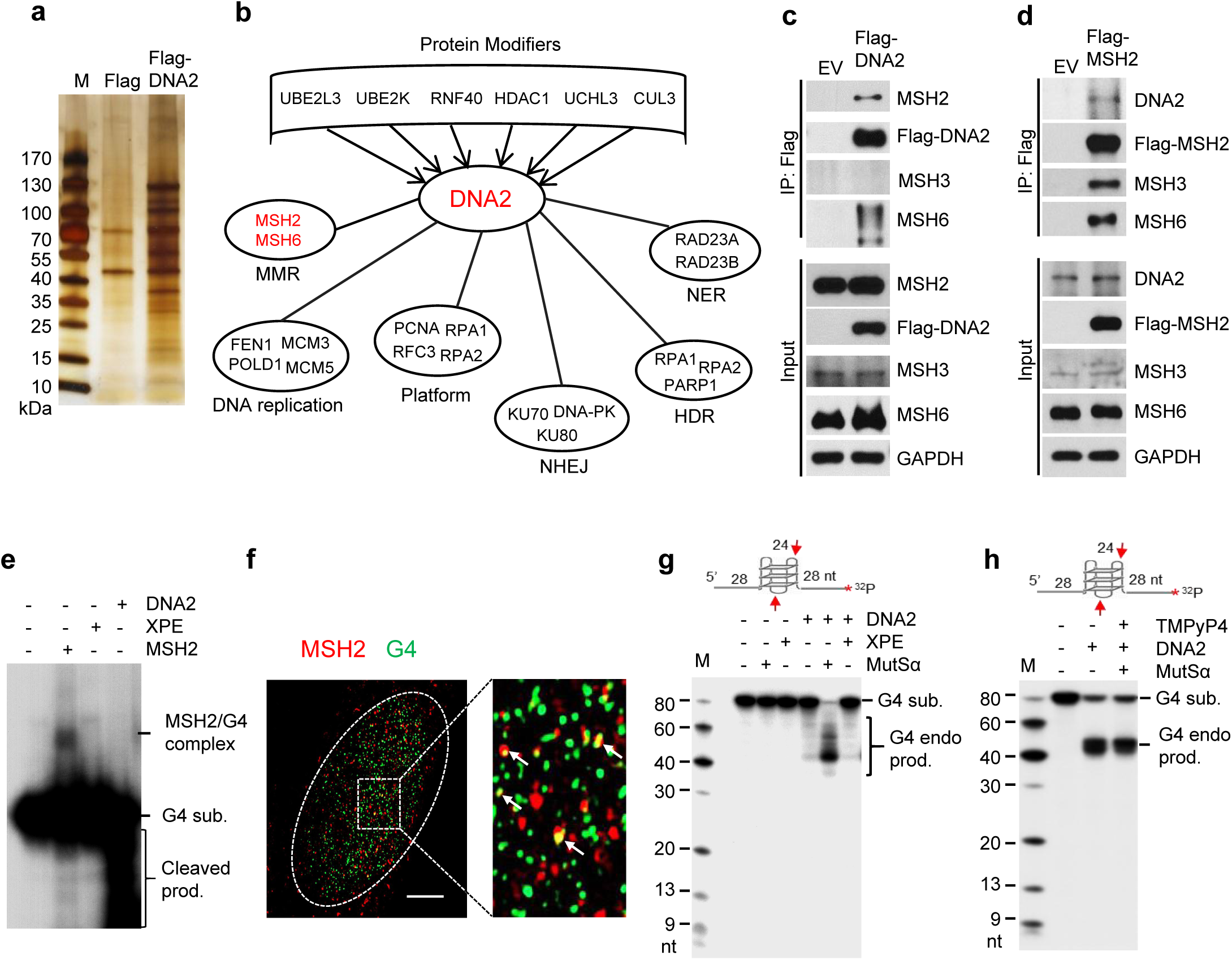
G4 binding protein complex MutSα stimulates DNA2-mediated G4 cleavage. **a** Silver staining SDS-PAGE showing the pulled-down proteins by Flag-tag M2 beads in whole cell extracts from 293T cells transfect with or empty vector of the vector encoding 3x-Flag human DNA2; **b** DNA replication and repair proteins co-pulled down with 3x-Flag-tagged DNA2; **c, d** Co-IP and western blot analysis of DNA2 interaction with MSH2 and MSH6; **e** The EMSA image shows the MutSα rather than XPE complex binds to the ^32^P-label G4 substrate. It also shows that DNA2 cleaves the G4 substrate; **f** AIRYSCAN confocal microscopy of G4s; MSH2-G4 co-immunofluorescence staining in MEFs. The right panel shows the enlarged view of the boxed region. The spots with co-localized MSH2 and G4 foci were indicated with white arrows. Scale bar = 5 μm; **g** The representative denaturing PAGE image shows DNA2 cleaving the ^32^P-labeled G4 substrate in the absence or presence of MutSα; **h** The representative denaturing PAGE image shows cleavage of the ^32^P-labeled G4 substrate by the DNA2-MutSα complex in the absence or presence of G4 stabilizer TMPyP4.

MSH2 and MSH6, which in complex is termed MutSα, recognize DNA mismatches, including those within DNA secondary structures ^37^. MutSβ, formed by MSH2 and MSH3, has recently shown to bind to G4 containing loop structures ^44^. Using co-IP and western blot analyses, we confirmed that MutSα components MSH2 and MSH6, but not MSH3 co-IPed with DNA2 (Fig. 2c) and that DNA2 co-IPed with MSH2 as well (Fig. 2d), demonstrating that DNA2 interacts with MutSα. Furthermore, gel-shift assays showed that MutSα could bind oligo-based G4 DNA substrates *in vitro* (Fig. 2e), and MSH2/G4 IF co-staining indicated that MSH2 foci colocalized with G4 foci in MEF cells (Fig. 2f). These findings suggest that MSH2 plays an important role in regulating the G4 excision function of DNA2. To determine if MutSα or MutSβ stimulates DNA2-mediated cleavage of G4s, we tested G4 cleavage by DNA2 in the absence and presence of MutSα or MutSβ. We found that MutSα, but not MutSβ, stimulated DNA2 to cleave G4 substrates *in vitro* (Fig. 2g, Supp. Fig. 3a). Surprisingly, MutSα also increased the unwinding activity of FANCJ in Polδ-driven primer extension on the G4 template, but MutSβ did not (Supp. Fig. 3b), indicating a potential role of MutSα in regulating multiple ways to resolve G4s. The NER component XPE, on the other hand, did not facilitate DNA2 to cleave G4 substrates (Fig. 2g). In addition, the G4 stabilizer TMPyP4, which was previously reported to block G4 unwinding by FANCJ ^45^, did not inhibit G4 excision by DNA2/MutSα (Fig. 2h). To determine the importance of MSH2 in G4 resolution, we analyzed G4 levels in WT and MSH2^-/-^ MEFs. Similar to DNA2 inhibition or deficiency, MSH2^-/-^ cells had significantly more G4 foci than the WT cells (Fig. 3a, b).

**Fig. 3.**
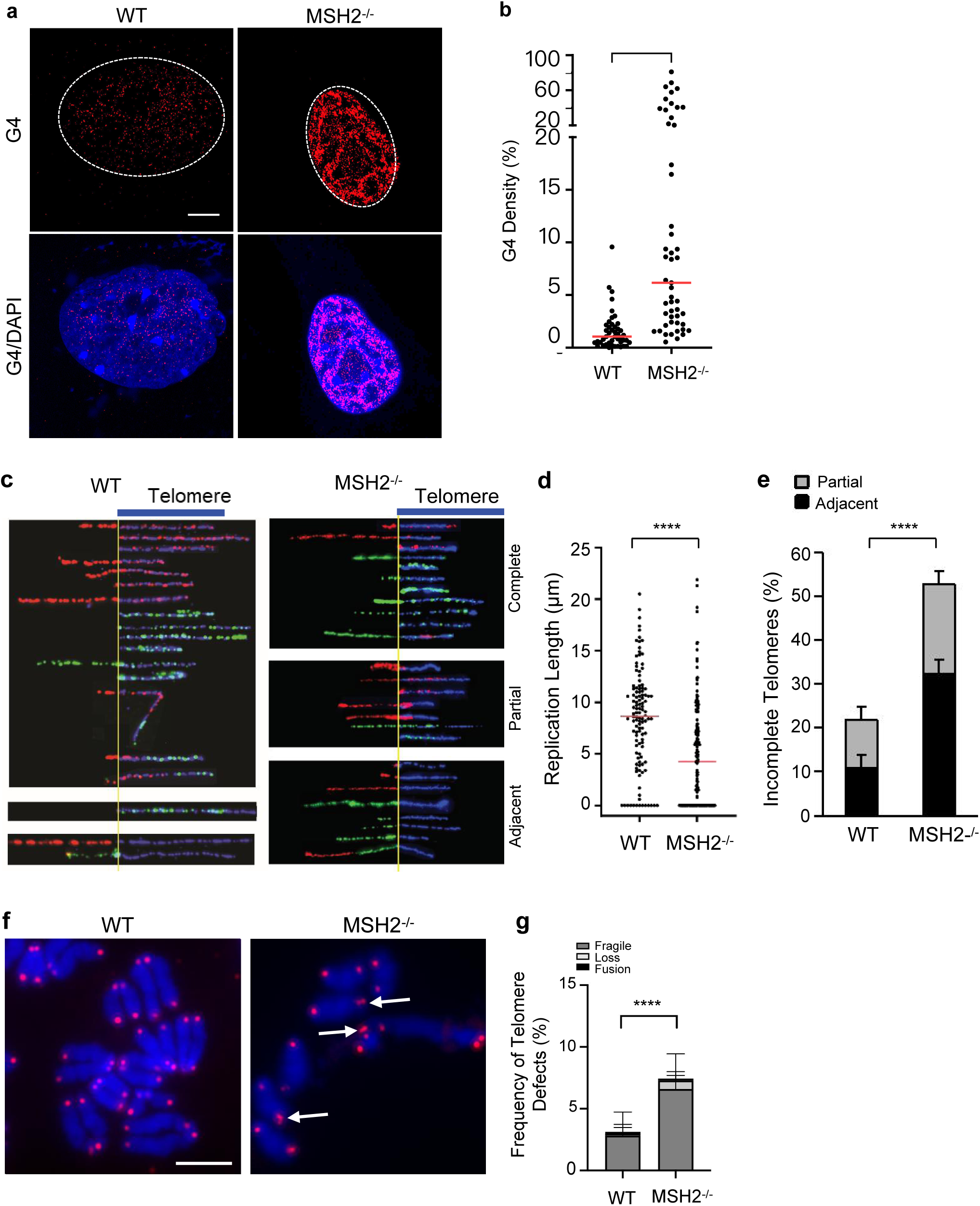
MSH2 deficiency results in G4 accumulation, defects in telomere replication. **a, b** G4 immunofluorescence staining in WT and MSH2^-/-^ MEFs. Panel A shows the representative AIRYSCAN confocal microscopy images of G4s, and Panel B shows the quantification of G4 in WT and MSH2^-/-^ MEFs. Scale bar = 5 μm; **c**-**e** SMARD analysis of WT and MSH2^-/-^ MEF cells. (**c**) Representative microscopy images of the replicated telomeres in WT or MSH2^-/-^; (**d**) Quantification of the relative length of replicated telomeres in different cells; and (**e**) Percentage of partial telomere replication and replication fork stalling adjacent to telomeres in WT or MSH2^-/-^ cells; **f** Telomere FISH images showing telomere abnormalities in the WT and MSH2^-/-^ MEFs. Scale bar = 5 μm; **g** Quantification of fragile telomeres in WT or MSH2^-/-^ cells treated with PIPER and capreomycin. All p values were calculated using the Two-way ANOVA. * p < 0.05, ** p < 0.01, *** p < 0.001, and **** p < 0.0001.

The findings that MutSα stimulates DNA2-mediated cleavage of G4s suggest that MutSα plays an important role in G4 excision to facilitate DNA replication at DTRs such as telomeres. To determine the biological significance of MSH2 in telomere replication, we carried out SMARD assays on WT and MSH2^-/-^ MEFs. Like DNA2^+/-^ MEFs, MSH2^-/-^ MEFs had significantly shorter replicated telomeres and showed more replication stalling at telomeres (Fig. 3c-e). On average, the length of replicated telomeres was 4.4 µm in MSH2^-/-^ MEFs, compared to 8.7 µm in the WT (Fig. 3d, 1f). 52% of telomeres showed fork stalling (32% partial and 20% adjacent) in MSH2^-/-^ MEFs, compared to ∼20% in the WT (Fig. 3e). Consistently, MSH2^-/-^ MEF cells had significantly more fragile telomeres (Fig. 3f, g).

### G4 stabilizing compounds block FANCJ-driven G4 unwinding, causing polymerase pausing, but do not affect DNA2-mediated G4 cleavage

Next, we sought to investigate the impact of environmental chemical compounds (ECCs) on G4 resolution in cells, particularly under a DNA2 haplo-insufficient or MSH2 knockout background. To identify possible G4-binding ECCs, we used our in-house-developed, structure-based, virtual ligand screening pipeline to screen a naturally occurring chemical library for potential binding to G4s (Supp. Fig. 4a). Potential G4-binding ECCs were defined as those with the highest docking scores (Supp. Table 1). From our docking screen, protoporphyrin IX (PPIX), a porphyrin derivative, and perylene derivatives pigment red 123 (PR123) (Supp. Fig. 4a) and PIPER (Supp. Fig. 4b) were among the ECCs with the highest potential for binding to G4. Consistently, PIPER and PPIX were both previously reported as potent G4-binding chemicals ^29, 46, 47^. In addition to the known G4-binding compounds, we found bleomycin, polyazo dye, aflatoxin B1, and capreomycin had high docking scores (model docking of ECCs to G4, Extended Data Fig. 4c). Therefore, we tested the impact of the known G4-binding perylene derivative PIPER, the porphyrin derivative TMPyP4, and the aminoglycoside antibiotic, capreomycin, on G4 resolution via helicase-driven G4 unwinding or DNA2-mediated G4 cleavage.

To determine the impact of these G4-stabilizing ECCs on G4 resolution via unwinding, we carried out Polδ-catalyzed primer extension on DNA templates which contained either a non-G4 forming sequence or a G4 forming sequence in the middle of the template (Supp. Fig. 5a). Polδ effectively extended the primer on the non-G4 forming DNA template, but it paused before the G4 forming sequence upon an increase in the K^+^ concentration, which is a G4-stabilizing ion, further confirming that this effect is due to the presence of G4 (Supp. Fig. 5b). Meanwhile, FANCJ, a typical helicase that unwinds G4s, relieved the pausing effect (Supp. Fig. 5c, d). Conversely, we found that the known G4-inducing/stabilizing compounds PIPER, TMPyP4, NNMP enhanced the pausing effect and reduced the fully extended product, but they did not affect Polδ extending on the non-G4 template (Fig. 4a and Supp. Fig. 6a, b). Consistent with our suggestion that capreomycin is a G4-binding ECC, we observed that it can also enhance Polδ pausing at the G4-forming sequence (Fig. 4b). We further found that PIPER inhibited FANCJ unwinding G4 (Fig. 4c). However, capreomycin did not remarkably inhibit FANCJ unwinding G4 (Fig. 4d). We observed that similar to TMPyP4 (Fig. 2h), neither PIPER nor capreomycin affected DNA2-mediated cleavage of the G4 DNA substrate (Fig. 4e, f). We further analyzed the impact of G4 stabilizing compounds on the accumulation of G4 foci in WT, DNA2^+/-^, and MSH2^-/-^ MEF cells. Treatment of WT MEFs with PIPER increased the accumulation of G4 foci compared to the untreated WT MEFs (Fig. 4g, h). DNA2 haploinsufficiency or MSH2 deficiency and 1 µM PIPER treatment potentiated each other in the accumulation of G4 foci. Capreomycin at 15 µM, like PIPER at 1 µM, caused an increase in G4 foci in WT MEFs and had a significant potentiation effect with DNA2 haploinsufficiency or MSH2 deficiency (Fig. 4g, h).

**Fig. 4.**
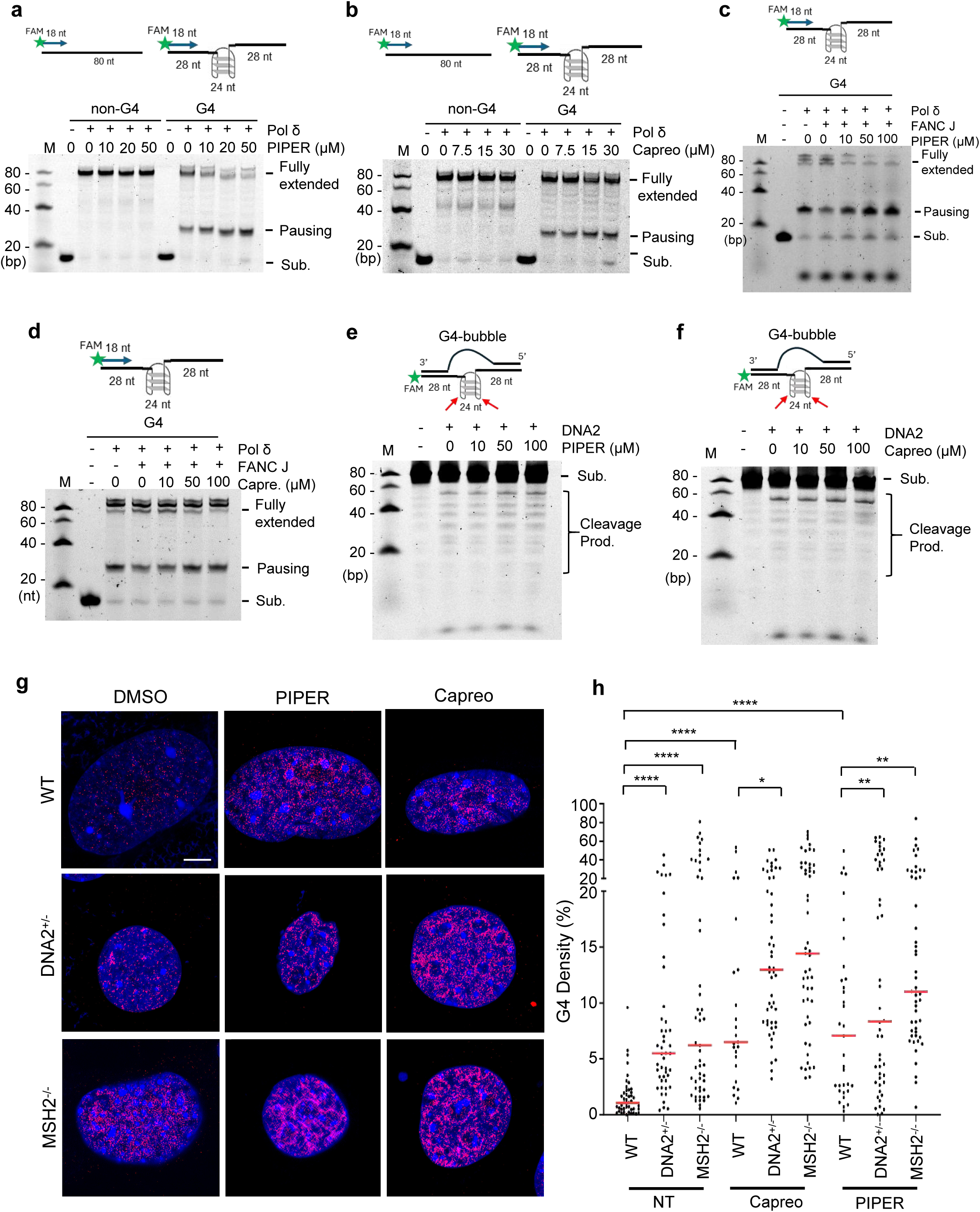
G4-stabilizing ECCs inhibit FANCJ unwinding G4, but not DNA2-mediated cleavage of G4 *in vitro*. **a, b** Primer extension on the non-G4 or G4 template in the presence of increasing concentrations of G4-stabilizing ECCs PIPER (**a**) and capreomycin (**b**). The top shows the diagram of the Polδ-catalyzed primer extension on a DNA template without or with a G4-forming sequence. The primer was labeled with FAM on the 5’ end. The bottom shows the representative denaturing-PAGE image of the primer extension assay; **c, d** Primer extension on the G4 template in the absence or presence of 5 nM FANCJ with or without 10, 50 and 100 μM PIPER (**c**) or capreomycin (**d**); **e, f** The cleavage of a FAM-labeled G4 substrate with increasing concentration of DNA2 in the absence or presence of increasing concentrations of PIPER (**e**) or capreomycin (**f**); **g, h** G4 immunofluorescence staining in WT and MSH2^-/-^ MEFs. (**g**) shows the representative AIRYSCAN confocal microscopy images of G4s, and (**h**) shows the quantification of G4 in WT, DNA2^+/-^, or MSH2^-/-^ MEFs without or with treatment with G4 stabilizers PIPER or capreomycin. Scale bar = 5 μm.

### G4-stabilizing compounds PIPER and capreomycin delay DNA replication at telomeres

To determine the impact of G4 accumulation caused by G4 stabilizing compounds on telomere replication in WT, DNA2^+/-^, and MSH2^-/-^ MEFs, we treated the cells with the G4-inducing/stabilizing compounds PIPER or capreomycin and performed SMARD to analyze replication dynamics at the telomeres of these MEFs. We found that both PIPER and capreomycin caused the slowing or stalling of replication at telomeres (Fig. 5a). Exposure to PIPER or capreomycin reduced the mean length of replicated telomeres from 8.7 µm (Fig. 1f) to 6.5 µm and 5.4 µm, respectively, in WT cells (Fig. 5b). Similarly, PIPER and capreomycin reduced the mean length of replicated telomeres from 4.2 µm (Fig. 1f) to 2.7 µm and 3.0 µm, respectively, in DNA2^+/-^ cells (Fig. 5b). However, PIPER and capreomycin did not cause a reduction in the mean length of replicated telomeres in MSH2^-/-^ MEFs (Fig. 3d, 5b). Meanwhile, we observed that PIPER and capreomycin increased the frequency of stalled replication forks at or in telomeres from 22% to 43% (Fig. 5c) in WT cells and increased the frequency from 50% to 55% and 59%, respectively, in DNA2^+/-^ cells (Fig. 5c). However, PIPER and capreomycin did not cause further fork stalling at the telomere in MSH2^-/-^ MEF cells (Fig. 3e, 5c).

**Fig. 5.**
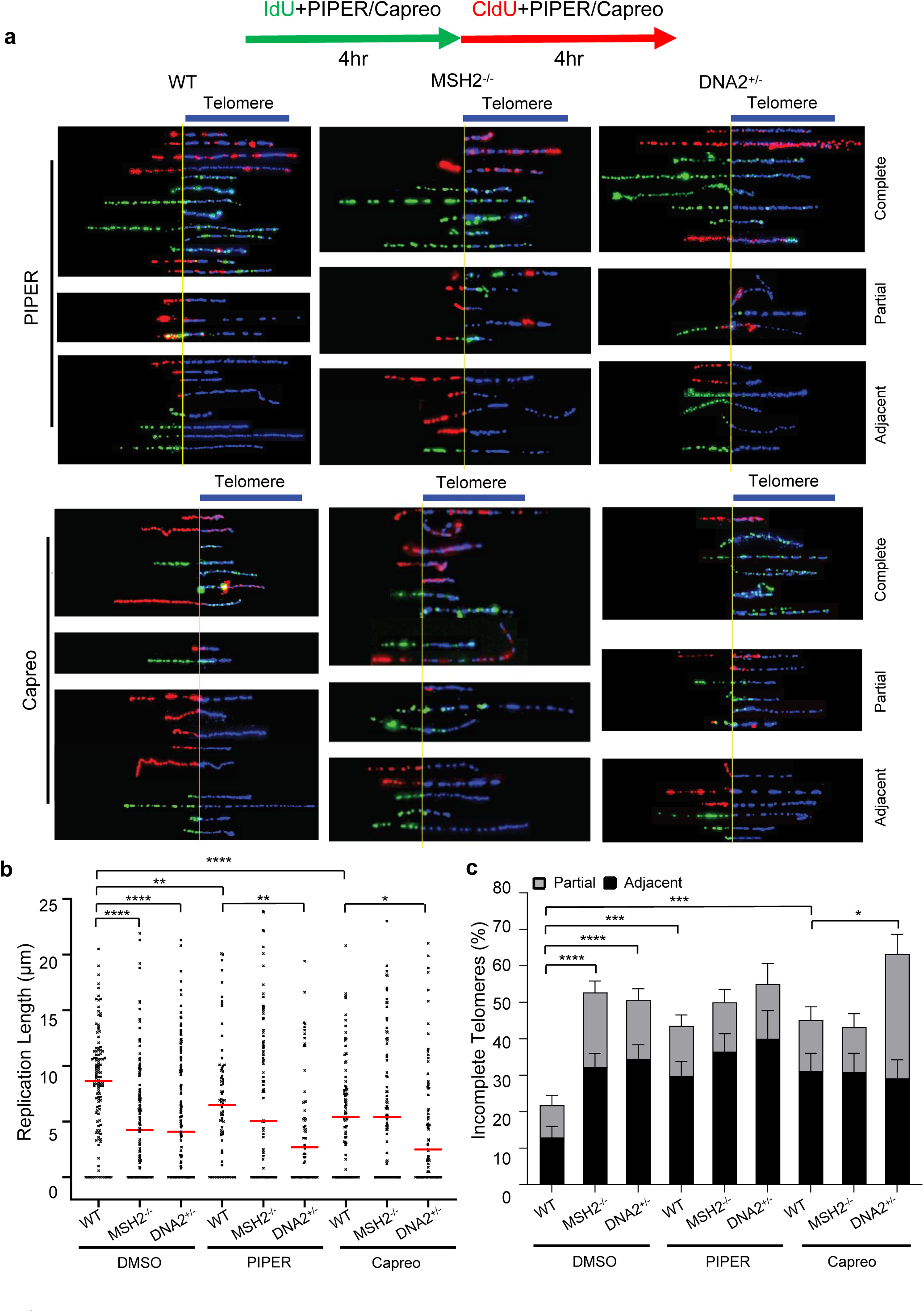
G4-stabilizing ECCs impair DNA replication at telomeres in WT DNA2^+/-^ and MSH2^-/-^ MEF cells. **a** SMARD microscopy images of replicated telomeres in WT, DNA2^+/-^, and MSH2^-/-^ MEF cells with or without exposure to PIPER or capreomycin; **b** Relative length of replicated telomeres in different cells; **c** Percentage of incompletely replicated telomeres in different cells. The frequency of partial telomere replication and replication fork stalling adjacent to telomeres was calculated.

Consistent with the result of telomere synthesis impairment due to exposure to PIPER and capreomycin, using telomere FISH we observed that PIPER or capreomycin exposure alone was sufficient to cause an increased frequency of telomere abnormalities including fragility, loss, and/or fusion in WT cells (Fig. 6a-d). DNA2^+/-^ MEF cells displayed significantly more telomere abnormalities than the WT MEF cells under normal culture condition, and exposure to PIPER or capreomycin caused additional telomere defects (Fig. 6a, b). Similar to DNA2^+/-^ MEF cells, MSH2^-/-^ MEF cells exhibited more sensitive to PIPER and capreomycin with a corresponding increase in telomere fragility, loss or fusion (Fig. 6c, d). It is worth noting that DNA2^+/-^ cells were more sensitive to PIPER or capreomycin than MSH2^-/-^ cells (Fig. 6a-d). It is possible that in the absence of MSH2, other proteins may facilitate DNA2 meditated G4 excision, albeit at much less efficiency.

**Fig. 6.**
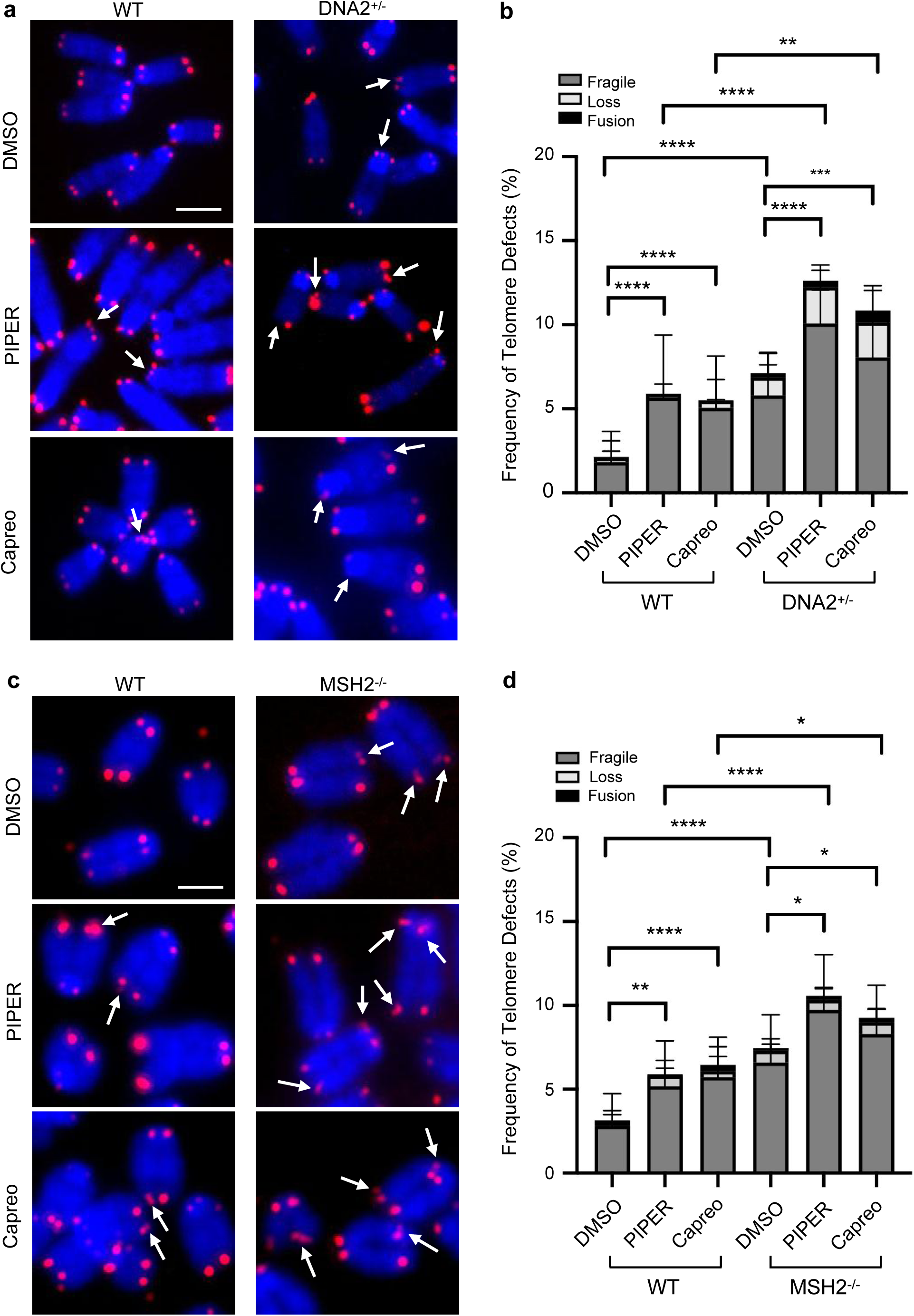
G4-stabilizing ECCs enhance telomere abnormalities in WT DNA2^+/-^ and MSH2^-/-^ MEF cells. **a** Telomere FISH images showing telomere abnormalities in the WT and DNA2^+/-^ MEFs treated with PIPER and capreomycin. Scale bar = 5 μm; **b** Quantification of fragile telomeres and shortened telomeres in different cells; **c** Telomere FISH images showing telomere abnormalities in the WT and MSH2^-/-^ MEFs treated with PIPER and capreomycin. Scale bar = 5 μm; **d** Quantification of fragile telomeres and shortened telomeres in different cells. All p values were calculated using the Two-way ANOVA. * p < 0.05, ** p < 0.01, *** p < 0.001, and **** p < 0.0001.

## Discussion

Our current studies define the biological importance of DNA2 nuclease mediated G4 excision. We show that DNA2 gene deficiency or treatment with a DNA2 inhibitor results in G4 accumulation in MEFs and various human cancer cells. We further determine the impact of G4-inducing/stabilizing chemicals on G4 excision as opposed to G4 unwinding, the well-studied G4 resolution mechanism. We show that FANCJ, a representative helicase that unwinds G4^45^, effectively assists Polδ in passing through the G4-forming sequence. However, G4-stabilizing ECCs, including PIPER and TMPyP4, inhibit FANCJ’s G4 unwinding activity, leading to Polδ passing at G4s. Meanwhile, G4 stabilization by PIPER or other G4-stabilizing ECCs has little effect on DNA2-mediated G4 excision. It suggests that two different G4 resolution mechanisms may function under different conditions. Under normal physiology, helicase driven G4 unwinding is the primary pathway, and G4 excision is an important pathway for efficient G4 resolution. However, DNA2-mediated G4 excision becomes the primary pathway for G4 resolution when cells are exposed to G4-stabilizing ECCs.

Given the nature of G-rich sequences to spontaneously form stable G4s or G4-like structures, G-rich sequences typified by telomere TTAGGG repeats are considered DTR regions. We previously used DNA2^+/-^ MEFs to show that DNA2 deficiency caused fragile telomeres ^31^, indirectly implicating DNA2 in telomere replication. In our current studies, using the single-molecule-based SMARD assay, we successfully assessed DNA replication at telomeres. We demonstrate that DNA2 deficiency or inhibition results in stalled replication forks at telomeres and telomere instabilities, especially in the presence of G4-stabilizing ECCs, including PIPER and a newly identified chemical stabilizer, capreomycin, in mammalian cells.

We further identify MutSα as a new G4-binding protein complex to facilitate G4 resolution via DNA2-mediated G4 excision and possibly G4 unwinding by FANCJ. We observe that MutSα binds G4 *in vitro,* and MSH2 localizes to G4 foci when G4s are induced by G4-stabilizing ECCs or G4 resolution is defective. Meanwhile, MSH2 interacts with DNA2 and stimulates DNA2 to cleave G4 structures. In addition to stimulating G4 excision, MutSα also facilitates FANCJ to unwind G4. Consistent with the biochemical data, MSH2 knockout, similar to DNA2 deficiency, causes G4 foci accumulation. Since G4 impedes replication fork progression ^48^, it is not surprising to see that proteins involved in G4 resolution are necessary to maintain replication dynamics through regions enriched in G4 structures (i.e., telomeres). As a result, cells likely require the use of alternative pathways to maintain telomeres, such as ALT telomere lengthening ^49^. Due to the problems in telomere replication, we show that cells lacking DNA2 are more prone to genome rearrangements, such as telomere fusion or sister chromatin exchange ^31^. We additionally show for the first time the role of environmentally contaminating compounds on telomere maintenance and the requirement for functioning DNA2 and MSH2 to resolve the impact of these compounds. While these compounds alone are sufficient to slow down normal replication of telomeres, the combination of DNA2 and/or MSH2 deficiency leads to a greater occurrence of fragile telomeres and other telomere defects.

In summary, our current studies provide several lines of evidence that DNA2-mediated G4 excision repair is important for resolution of G4 in mammalian cells, especially in the presence of G4 stabilizing compounds, which suppress G4 unwinding by helicases. Furthermore, we define the DNA repair protein complex MutSα that recognizes G4s, recruits DNA2, and stimulates its cleavage of G4s. G4 accumulation causes replication stress, which is a major source of genome instability. We demonstrate that DNA2 or MSH2 deficiency in MEF cells causes telomere instability. Therefore, DNA2 haploinsufficiency that causes defective G4 excision leads to genome instability and contributes to tumorigenesis ^31^. On the other hand, the results from our current studies also have implication in cancer cell survival in response to anti-cancer treatment. G4 induction and stabilization by drugs such as CX5641 have been used as a strategy to create overwhelming replication stress to kill cancer cells ^50^. Robust G4 excision isa hurdle for the effective killing of cancer cells by G4-stabilizing drugs. Simultaneous induction of G4 formation and inhibition of G4 excision could be an effective combination for future cancer therapy.

## Methods

### DNA2 Pulldown and Mass Spectrometry

3x-Flag-tagged DNA2 was expressed or pulled down with anti-Flag tag M2 beads as previously described ^31, 42, 51^. Briefly, cells transfected with the empty vector (control) or the vector encoding 3x-Flag-DNA2 were lysed by brief sonication in an immunoprecipitation (IP) buffer containing 50 mM HEPES-KOH (pH7.4), 150 mM NaCl, 0.1% NP40, 10% glycerol, and 1x protein inhibitor cocktail (ThermoFisher). After centrifugation (20,000 g, 15 min, 4°C), the clear supernatant was incubated with anti-Flag M2 magnetic beads (Sigma) overnight. The beads were washed with a wash buffer containing 50 mM HEPES-KOH (pH 7.4), 500 mM NaCl, 0.1% NP40 and 10% glycerol, and 1x protein inhibitor cocktail once and the IP buffer once. The DNA2 protein complexes were eluted with 3x-Flag peptide (250 μg/ml). The eluted protein complexes and the control were run into a 4-15% Mini-PROTEAN® TGX™ Precast Protein Gel (Bio-Rad) and stained with a silver staining kit or Coomassie brilliant blue (ThermoFisher Scientific). All protein bands were de-stained and excised, followed by in-gel digestion, extraction, and LC/MS analysis using a Thermo Scientific Orbitrap Fusion Mass Spectrometer following the supplier’s instruction. The proteins were identified using both Proteome Discoverer Software with Sequest (Version 2.0) and the Mascot algorithm (Mascot 2.5.1). The mass spectrometry proteomics data have been deposited to the ProteomeXchange Consortium via the PRIDE partner repository with the dataset identifier “PXD059843”.

### Immunoprecipitation and western blot

Immunoprecipitation (IP) was carried out as previously described ^51^. Briefly, cells were lysed in a buffer containing 0.5% Nonidet P-40, 50 mM Tris-HCl, 0.1 mM EDTA, 150 mM NaCl, and proteinase inhibitors. Whole-cell lysates were incubated with the antibody against a specific protein and Pierce^TM^ Protein A/G Magnetic Beads (ThermoFisher Scientific). The beads were washed with a wash buffer containing 0.5% Nonidet P-40, 50 mM Tris-HCl, 0.1 mM EDTA, 500 mM NaCl, and proteinase inhibitors. The IPed proteins were eluted with 2x SDS loading buffer and analyzed by western blot following the standard western blot protocol.

### DNA2 Nuclease Assay

To make the G4 substrate, the G4-B oligo was labeled with FAM at the 3′ end. The oligo sequences are specified in Supplementary Table S2. The G4 structure substrate was made according to a previously described protocol ^31^. The formation of G4, which migrates faster than the corresponding non-G4, was confirmed by non-denaturing PAGE (Supp Fig. S7). To assay the nuclease activities, 40 fmol (5 ng), 80 fmol (10 ng) and 160 fmol (20 ng) DNA2 protein was incubated with 50 nM DNA substrate in a reaction buffer containing 20 mM Tris-HCl (pH 7.5), 8 mM MgCl_2_, 1 mM DTT, 10 mM KCl, 2 mM ATP and 0.1 mg/ml BSA at 30°C for 30 minutes. The DNA substrates and products were analyzed with a 15% denaturing PAGE gel.

### G4 Staining in Cell Culture and Foci Quantification

Cells were fixed using 4% PFA in PBS for 15 min at RT, followed by 0.5% Triton X-100 (15 min, RT), and blocking in 5% BSA + 0.1% Tween-20 and Image iT-FX Signal Enhancer. The anti-G4 primary antibody (MABE1126, Sigma) was diluted 1:500 in 4% BSA and 0.1% Tween-20 overnight at 4°C, washed with 0.1% Tween-20, and stained with 1:1000 GAM Alexa Fluor 568 for 1 hr at RT. Cells were counterstained with DAPI and mounted in ProLong Gold (Thermo).

Imaging was performed using the Zeiss LSM 900 with Airyscan 2. Briefly, we zoomed into individual nuclei and scanned with a resolution such that a theoretical point spread function would be sampled at least 3x. After acquisition, images were processed using Zeiss’s Airyscan Joint Deconvolution algorithm at 10 iterations to ensure maximum resolution.

To quantify foci, we used the particle finder algorithm on ImageJ. We used a consistent threshold across all imaging conditions and counted the number of foci over at least 15 cells per condition.

### Single Molecule Analysis of Replicated DNA

SMARD was carried out as previously described ^36^. Cells were grown in DMEM with 10% FBS (Gibco) and pen/strep (Gen Clone). Subconfluent, dividing cells were successively pulse-labeled with IdU (30 μM, 4 hr, Sigma) and then CldU (30 μM, 4 hr, Sigma). After labeling, cells were trypsinized, harvested, and cast into 0.5% agarose plugs (TopVision). Cells were lysed at 50°C in 1% n-lauroylsarcosine, 0.5 M EDTA, and 0.2 mg/ml Proteinase K (Bioland). DNA was digested outside of telomere regions using *SwaI* (NEB) overnight at RT. Digested DNA was then separated by electrophoresis with 0.7% SeaPlaque GTG Agarose (Lonza) in 0.5X TBE buffer. DNA fragments from 15-50 kb were excised and the agarose digested with β-agarase (NEB). This DNA was then stretched on 3-aminopropyltriethoxysilane-treated coverslips (Sigma), denatured with NaOH in 70% ethanol and fixed with glutaraldehyde. DNA was hybridized with the TelC biotinylated telomere probe (PNA Bio) at 37°C for 2 hours. After hybridization, DNA was blocked with 5% BSA and stained with Avidin Alexa Fluor 350 (Thermo), Mouse Anti-BrdU (BD) specific for IdU, and Rat Anti-BrdU (Abcam) specific for CldU. DNA was then stained again with a Goat Anti-Mouse Alexa 568 (Invitrogen), Goat Anti-Rat DyLight 488 (Invitrogen), and biotinylated Goat Anti-Avidin D (Vector Labs). Alternating rounds of avidin and anti-avidin were used to amplify the signal of the FISH probe. Coverslips were mounted on slides using ProLong Gold (Thermo) and imaged using the Zeiss Observer II with a 63x objective. Images were processed using ImageJ and aligned using Adobe Illustrator.

### Telomere Fluorescence In Situ Hybridization

FISH on telomeres were performed as described previously ^31^. For FISH, cells were treated with colcemid, collected by trypsinization, swollen in 0.075 M KCl at 37°C for 15 min, and fixed in a freshly prepared 3:1 mix of methanol:glacial acetic acid. Slides containing metaphase spreads were then denatured and hybridized to peptide nucleic acid (PNA) probe Cy3-OO-(CCCTAA)3 to visualize telomeres. Slides were then washed and dehydrated in ethanol series. DNA was counterstained with DAPI, and images were taken under Zeiss Observer epifluorescence microscope with a 100× objective.

### Polδ Extension Assay

FAM-labeled primer was annealed to the G4-B oligo and the G4 structure was formed in the G4 folding buffer containing 10 mM Tris-HCl (pH 7.5), 50 mM KCl at 95°C for 5 minutes, 72°C for 10 minutes, and then cooled down to room temperature. To perform the polymerase extension assay, 1.98 pmol (500 ng) Polδ was incubated with 50 nM DNA substrates in an extension buffer (20 mM Tris-HCl (pH 7.5), 8 mM MgCl_2_, 1 mM DTT, 0.10 mg/ml BSA, 10 mM KCl, 1 mM ATP, 250 μM dNTP) with or without the indicated amounts of FANCJ at 30°C for 30 minutes. The primer extension reaction was stopped with an equal volume of a 2x STOP solution (10 mM EDTA in formamide) at 95°C for 15 minutes. Subsequently, the extensions were resolved on a denaturing PA-urea gel (1x TBE, 7 M urea, 15% PAA, Acrylamide/ Bisacrylamide ratio 19:1) in 1x TBE run at 100V for 1 hour 20 minutes. Gels were imaged with a Typhoon FLA9500 scanner (GE Healthcare) and quantified with ImageJ. For the polymerase extension assay with G4 stabilizers, TMPyP4, PIPER, NMMP, and capreomycin, 50 nM DNA substrate was incubated with the indicated amounts of G4 stabilizers in the extension buffer at room temperature for 5 minutes, and then the binding products were performed for primer extension assay.

### High-Throughput Virtual Screening to Identify G4 Stabilizing Compounds

We employed a high-throughput virtual screening (HTVS) of 10,449 environmental contaminants to identify potential high-affinity G4 stabilizers. Compound structures were curated from the US Environmental Protection Agency (EPA) CRD2016 database (8,708 chemicals from USA industrial and commercial usage), EPA hazardous waste list (724 chemicals), Spin database (1,229 industrial chemicals from Nordic countries), Keml database (536 restricted chemicals in Sweden), EPA CCL4 chemicals (96 water-contaminated chemicals), FDA-approved drugs or compounds in clinical studies (6,605 molecules from DrugBank), and food toxins and metabolites (210 compounds). Additionally, 36 previously reported G4 stabilizers from the literature and Protein Data Bank (PDB), 37 close structural analogs (similarity cutoff: 0.7), 22 aflatoxin family compounds, and 188 perylene family compounds were included in the analysis. The curated library was docked against G4 crystal structure PDB: 3UYH. Prior to the molecular docking analysis, the curated library and G-quadruplex PDB structure were preprocessed using the LigPrep and Protein Preparation wizards, respectively, within the Maestro docking software (Schrödinger) to resolve common structural issues and adjust the pH to physiological conditions (pH 7.4 ± 1.0).

Potential G4 stabilizers were identified by comparing the curated compounds’ docking scores, chemical structures, and binding conformations to the 36 previously reported G4 stabilizers. Docking scores, shown as ΔG (free energy change), have been provided for the top nine potential G4-binding ECCs identified from HTVS (Supplementary Table S1). The most tractable G4 stabilizer candidates were subsequently validated *in vivo*, leading to the selection of PIPER and capreomycin as potent representative G4 stabilizing compounds for the *in vitro* experiments within this study.

## Acknowledgements

We thank Dr. Brian Armstrong at the COH Light Microscopy core facility for his technical support of AIRYSCAN confocal microscopy and SMARD analysis. This work was supported by NIH grants R50 CA211397 to L.Z. and R01 CA085344 to B.H.S. Research reported in this publication includes work performed using City of Hope shared resources supported by the National Cancer Institute of the National Institutes of Health under award number P30 CA033572. This research was also supported in part by the Intramural Research Program of the NIH, National Institute on Aging.

## Author Contributions

A.F., T.Z., C.S., H.L., H.Y., N.L., M.Z., and G.S., conducted biochemical and cellular experiments. S.E. conducted *in vivo* experiments. J.H. did the virtual ECC screening for G4 stabilizers. J.S. and R.B. provided purified recombinant FANCJ protein and contributed to the design of helicase-relevant experiments. V.G. and W.C. helped with telomere FISH experiments and provided technical advice on data analysis. S.K, D.Z., and C.S. provided SMARD protocols and technical advice. W.E. provided MSH2^-/-^ MEF cells. G.M.L. provided the purified MutSα protein complex. M.Y.W.T. Lee provided the monoclonal antibody used to purify the recombinant DNA polymerase delta protein complex. L.Z. designed biochemical and cellular experiments, analyzed the data, and wrote the manuscript. B.S. supervised the entire project, designed and coordinated the majority of experiments, provided input, and finalized the manuscript.

## Conflict of Interest

The authors declare no conflict of interest.

## Notes

### Competing Interest Statement

The authors have declared no competing interest.

