## Supplementary Tables and Figures for "DNA2 and MSH2 activity collectively mediate chemically stabilized G4 for efficient telomere replication"

### Supplementary Information

**Supplementary Table 1: List of top ECCs that dock to the G4 structure**

| CAS NO. | Common Name | Category | Docking Score (kcal/mol) |
|---|---|---|---|
| 11056-06-7 | Bleomycin | FDA drug | -14.871 |
| 213416-70-7 | PIPER* | Small mol. | -10.544 |
| 24106-89-2 | Pigment Red 123* | CDR comp. | -9.145 |
| 67786-25-8 | Stilbenedisulfonate | CDR comp. | -7.326 |
| 1162-65-8 | Aflatoxin B1 | Carcinogen | -6.365 |
| 112484-44-3 | Polyaza dye | CDR comp. | -6.014 |
| 553-12-8 | Protoporphyrin Ⅸ | Metabolite | -5.076 |
| 11003-38-6 | Capreomycin | FDA drug | -5.026 |
| 146939-27-7 | Ziprasidone | FDA drug | -4.476 |

*Perylene derivatives

**Supplementary Table 2: List of oligonucleotides for DNA2 nuclease assay and Polδ extension assay**

| oligonucleotide name | sequence (5' to 3') |
|---|---|
| G4B/FAM-G4B | GTTAAGATAGGTCTGCTTGGCATGTCAATTAGG GTTAGGGTTAGGGTTAGGGCTCTGTGGTTGAG GCAGAGTCCTTAAGC |
| Complemented-G4B | GCTTAAGGACTCTGCCTCAACCACAGAGCCCT AACCCTAACCCTAACCCTAATTGACATGCCAAG CAGACCTATCTTAAC |
| Random/FAM-random | GTTAAGATAGGTCTGCTTGGCATGTCAAGGTTT CTAAAGAAGCCGACGGTAGCTCTGTGGTTGAG GCAGAGTCCTTAAG |
| FAM-G4B-primer | GCTTAAGGACTCTGCC |
| G4B (blocking 3'-end cleavage) | TGCTCGTTTTGTTTGGTCTGCTTGGCATGTCAA TTAGGGTTAGGGTTAGGGTTAGGGCTCTGTGG TTGAGGCAGAGTCCTTAAGC |
| Complemented-G4B (blocking 3'-end cleavage) | TGCTCGCTTTTGGTCTCTGCCTCAACCACAGAG CCCTAACCCTAACCCTAACCCTAATTGACATGC CAAGCAGACCTATCTTAAC |

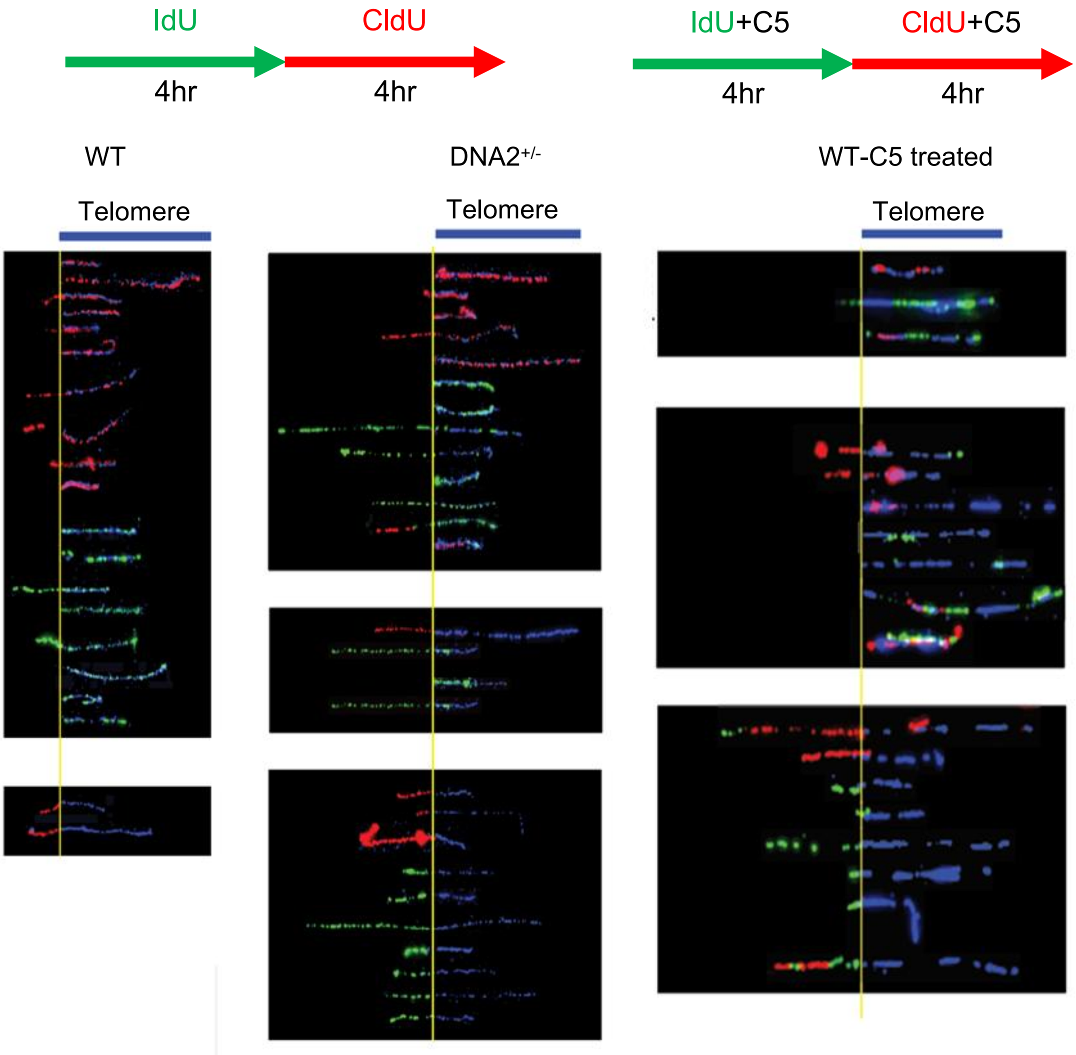


**Supplementary Fig. 1 | SMARD assays on MEF cells of WT, DNA2$^{+/-}$ and WT treated with C5.** Top: Scheme of the IdU (green) and CldU (red) pulse labeling and C5 treatment. Bottom: Representative images for SMARD assay results. WT and DNA2$^{+/-}$ MEF cells or WT MEF cells treated with C5 were labeled with IdU/CldU. DNA was digested with a cutting enzyme and isolated. Telomeric DNA was identified by a TelC-Biotin probe and fluorescently labeled Avidin (blue). Replicating DNA, which was incorporated with IdU (green) and/or CldU (red), was detected using anti-BrdU antibodies and Alexa-488 or Alexa-568 conjugated secondary antibodies.

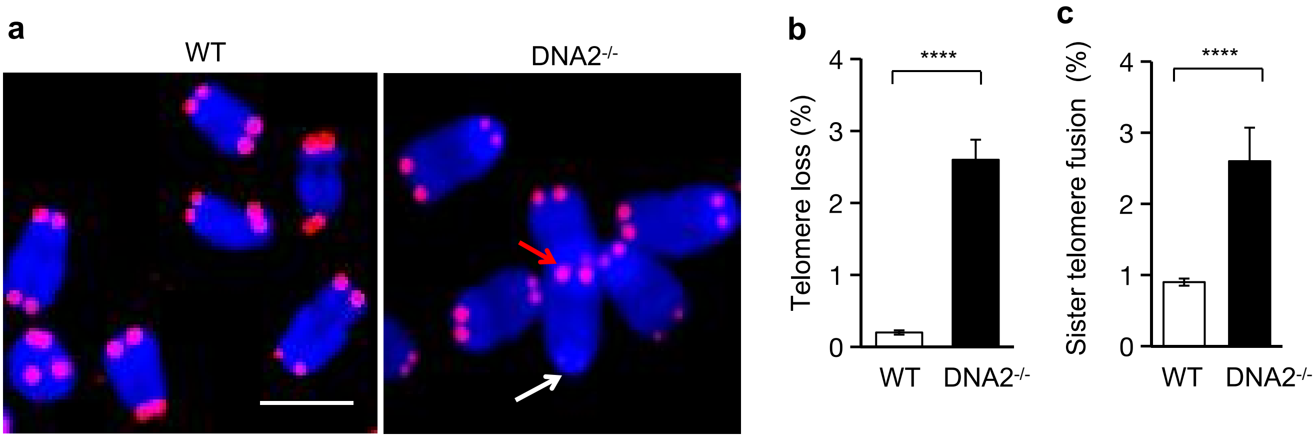


**Supplementary Fig. 2 | Telomere stability in WT and DNA2$^{-/-}$ mouse ES cells. a** Representative telomere FISH images showing telomeres in WT and DNA2$^{-/-}$ mouse ES cells. DNA was counter-stained by DAPI (blue). Telomere loss (signal-free ends) or sister telomere fusion are indicated by white and red arrows, respectively. Scale bar = 5 μm; **b, c** Quantification of telomere loss and sister telomere fusion in mouse ES cells. Values are mean ± st.d. of four assays.

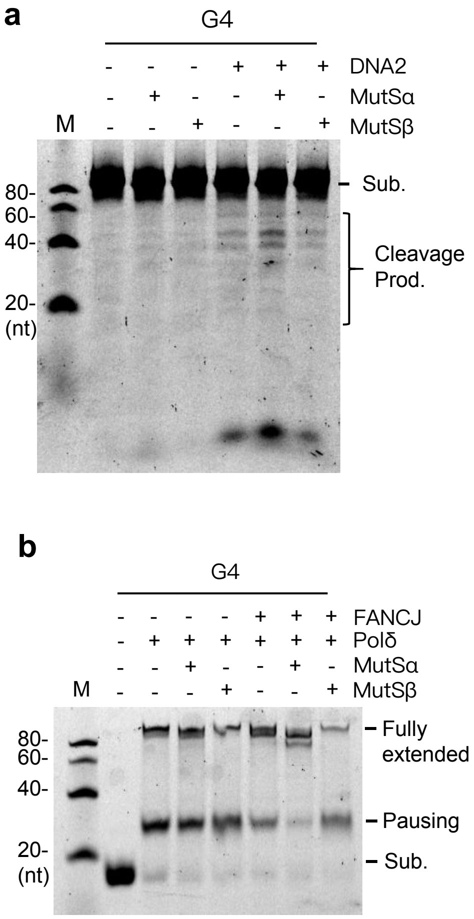


**Supplementary Fig. 3 | MutSα but not MutSβ stimulates G4 cleavage by DNA2 or G4 unwinding by FANCJ. a** The cleavage of a FAM-labeled G4 substrate by DNA2 in the absence or presence of MutSα or MutSβ; **b** FAM-G4B-primer extension on the G4 containing DNA template by Polδ in the absence or presence of FANCJ alone or in combination with MutSα or MutSβ.

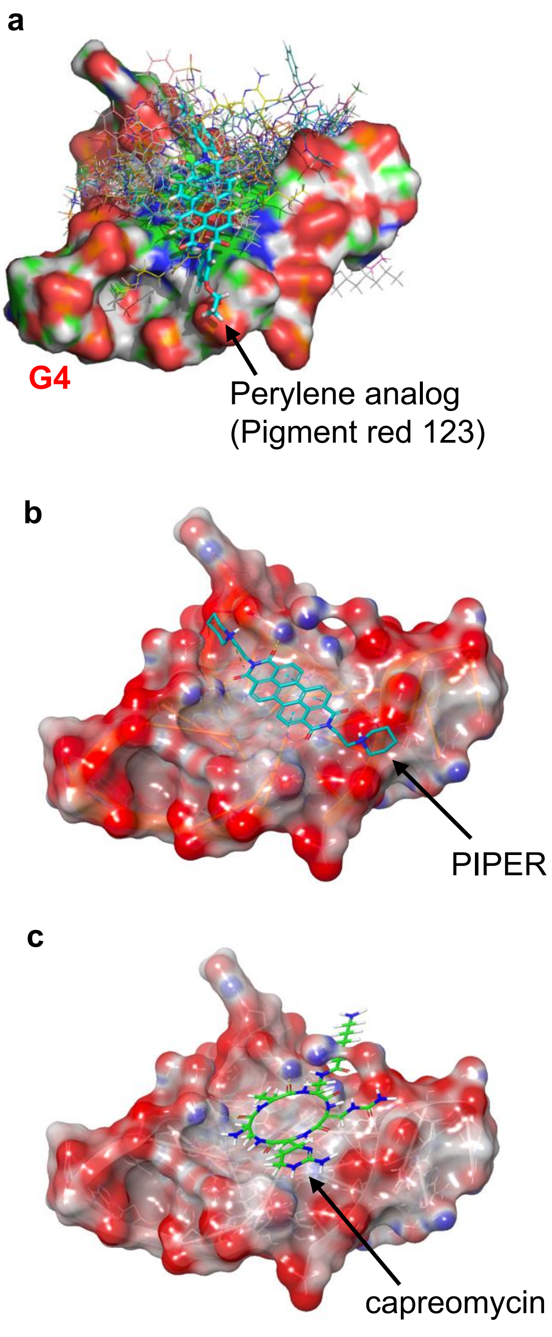


**Supplementary Fig. 4 | Virtual screening to identify potential ECCs that bind to G4. a** Docking different ECCs onto a G4 structure (PDB 3uyh). Docking of a known G4 binding compound, perylene analog Pigment Red 123 is specified; **b, c** Modeling of the known G4 binding ECC PIPER (**b**) and a candidate G4-binding ECC, capreomycin (**c**), onto the G4 structure. The aromatic region of the PIPER molecule (light blue) (**b**) and the capreomycin molecule (green) (**c**) extend diagonally across the planar surface of the tetrad. The side chains of PIPER are embedded in the grooves (**b**), while the capreomycin molecule shows slight favoring towards the 5' end of the G4 (**c**). The electron density surface for the 3uyh structure is shown and depicts the regions of most positive potential (dark blue) and most negative potential (dark red).

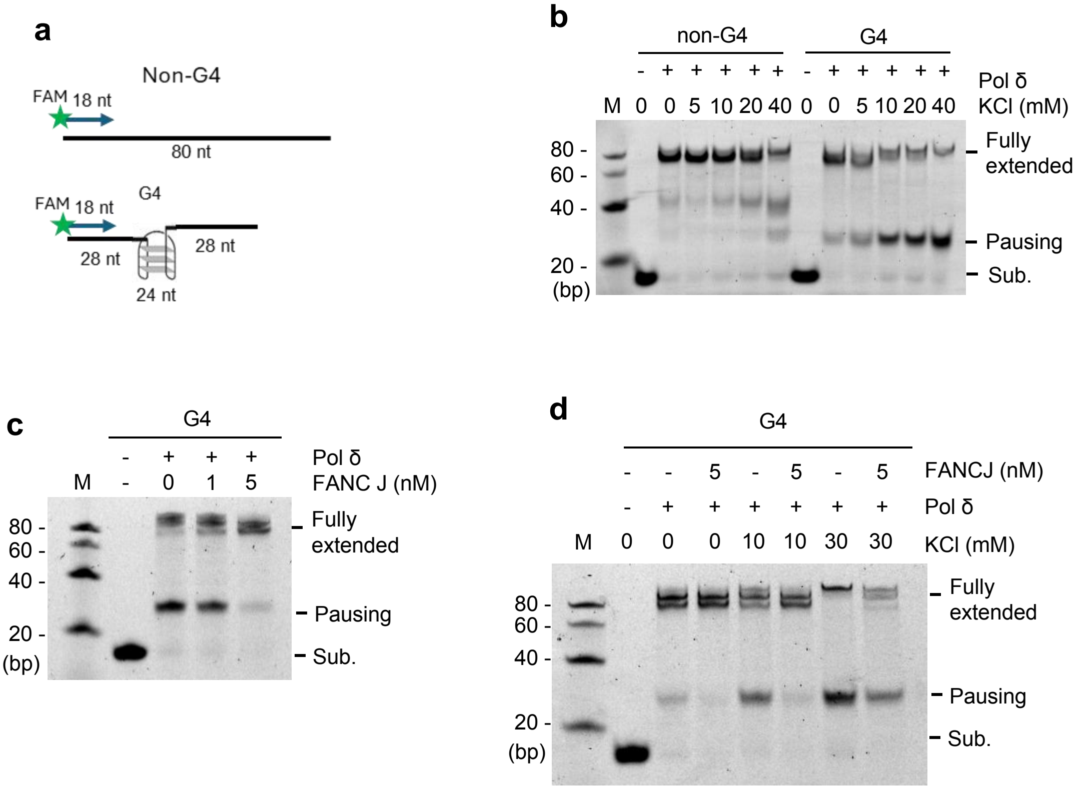


**Supplementary Fig. 5 | Primer extension on the non-G4 and G4 DNA template. a** The diagram shows the Polδ-catalyzed primer extension on the template without G4 (non-G4) or with G4 (G4) forming sequence. The primer was labeled with FAM on the 5' end; **b** Polδ-catalyzed primer extension on the template of non-G4 and G4 in the presence of increased concentration of KCl; **c** Polδ-catalyzed primer extension on the template of G4 with different concentrations of FANCJ; **d** Polδ-catalyzed primer extension on the template of G4 with FANCJ in the presence of different concentrations of KCl.

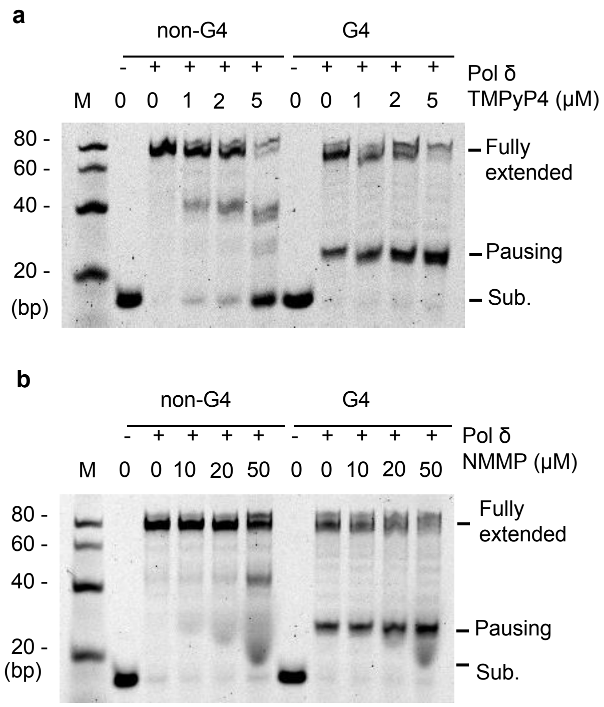


**Supplementary Fig. 6 | Impact of G4-stabilizing compounds TMPyP4 and NMMP on primer extension on the non-G4 and G4 DNA template. a, b** Polδ-catalyzed primer extension on the template of non-G4 and G4 in the presence of increasing concentrations of TMPyP4 (**a**) and NMMP (**b**).

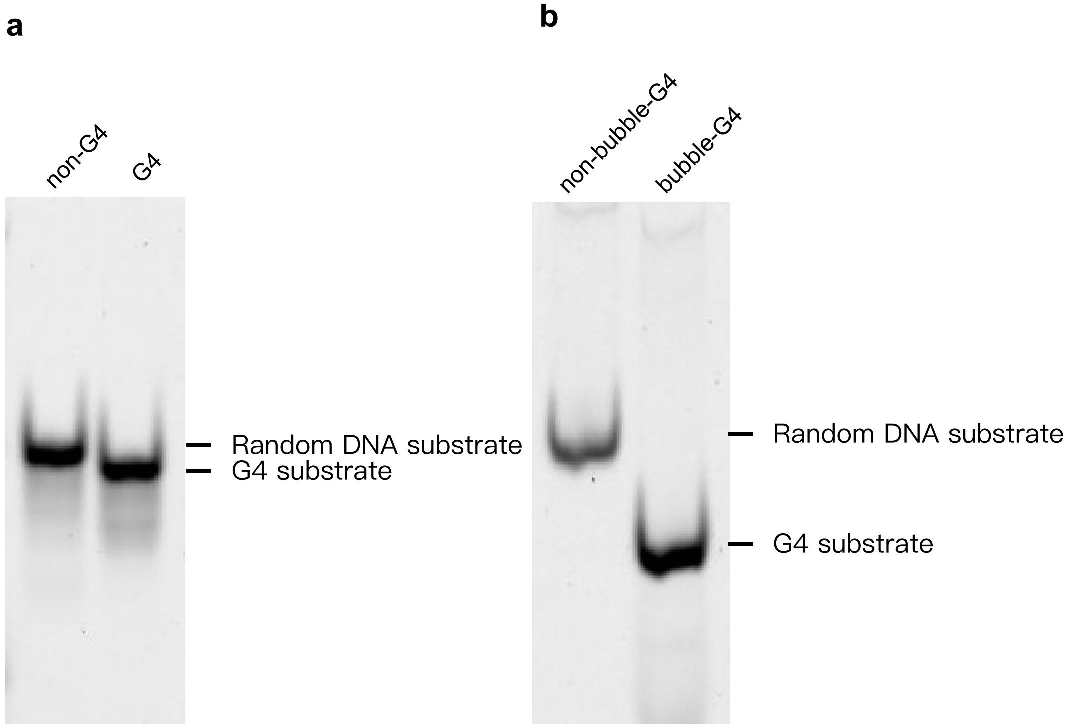


**Supplementary Fig. 7 | Native PAGE confirms the formation of G4 DNA substrates.** **a** Single-stranded DNA substrates were annealed using an oligo of random DNA sequence or an oligo containing a G4-forming sequence. The formation of G4 was analyzed using 8% native PAGE; **b** DNA bubble substrates were prepared with the oligo of random sequence or the oligo containing a G4-forming sequence. The formation of G4 was analyzed using 8% native PAGE.
